# PKD-1 Signaling Is Required for the Maintenance of CSCs with Epithelial-mesenchymal Plasticity in Pancreatic Neuroendocrine Tumors

**DOI:** 10.1101/2022.02.17.480869

**Authors:** Yichen Guo, Yinan Jiang, J. Bart Rose, Renata Jaskula-Sztul, Anita B. Hjelmeland, Herbert Chen, Bin Ren

## Abstract

Pancreatic neuroendocrine tumors (pNETs) are extremely heterogeneous and highly vascularized neoplasms that arise from endocrine cells in the pancreas. pNETs harbor a subpopulation of stem cell-like cancer cells (cancer stem cells/CSCs), which contribute to intratumoral heterogeneity and promote tumor maintenance and recurrence. In this study we demonstrated that CSCs in human pNETs co-expressed PKD-1 and CD44. We further identified PKD-1 signaling as a critical pathway in the regulation of CSC maintenance in pNET cells. PKD-1 signaling regulated the expression of a CSC-related gene signature and promoted CSC self-renewal. Intriguingly, pharmacological or genetic disruption of PKD-1 signaling in human pNET cells impaired lysophosphatidic acid-induced expression of genes associated with epithelial to mesenchymal transition (EMT), specifically E-cadherin and vimentin. This study indicates that PKD-1 signaling is essential for the maintenance of a subpopulation of CSCs in pNETs at an intermediate state along the epithelial–mesenchymal spectrum, thereby leading to a CSC phenotype with plasticity and partial EMT. Inhibiting the PKD-1 pathway may facilitate the elimination of CSC subpopulations to curb pNET progression, therapeutic resistance and metastasis. Given that the signaling networks associated with CSC maintenance and EMT are complex and extend across multiple levels of gene regulation, this study provides insight into signaling regulation of partial EMT in determining CSC fate and aids in developing potential therapeutic strategies that target different subsets of CSCs in a variety of cancers.

## 1. Introduction

Pancreatic neuroendocrine tumors (pNETs) are a group of heterogeneous, extremely vascularized neoplasms that arise from endocrine cells in the pancreas. Upon diagnosis, 60-70% of patients have a metastatic disease [1] and are no longer candidates for surgical intervention. The highly vascularized pNETs are characterized by elevated levels of vascular endothelial growth factor (VEGF) and its receptors [2], which are key factors in the regulation of angiogenesis [3,4]. Targeted therapy against the VEGF signaling pathway has been approved in the treatment of advanced pNETs including unresectable and metastatic disease. However, the therapeutic efficacy of antiangiogenic therapy is limited, due, in part, to the potential for increased invasion and an association with high rates of metastases [5,6].

Considering for the intratumoral heterogeneity of cancer cells associated with the development of cancer stem-like cells (CSCs) and potential value of targeting the vascular niches in CSCs [7-10], understanding mechanisms by which CSCs are regulated will provide insight into the development of potential anti-cancer therapies. CSCs are able to self-renew, have higher tumorigenic potential, and generate heterogeneous lineages of cancer cells, thereby promoting neoplastic maintenance, heterogeneity, and metastasis [7,8,11,12]. These CSCs have been identified in a number of solid tumors, including pNETs [13,14]. Though critical to the development of novel therapeutic approaches to advanced pNETs, understanding the mechanisms of CSC maintenance in pNETs has proven challenging but is critical for developing novel therapeutic approaches against advanced pNETs.

Many signaling pathways including chromatin remodeling, Notch1 and PI3K/AKT/mTOR signaling are central to the genetic heterogeneity of pNETs. These pathways may be critical for the regulation of CSC properties and pNET progression [6,8,15,16]. However, little is known about the signaling mechanisms by which pNET CSCs are regulated during tumor maintenance or progression. Protein kinase D (PKD-1), a member of the serine/threonine kinase D family, is involved in the regulation of chromatin remodeling by modulating histone deacetylases [17,18]. PKD-1 also activates PI3K/Akt signaling and regulates Notch 1 signaling in several different cell types, including vascular endothelial cells and cancer cells [3,19-25]. Among many biological outcomes of PKD-1 signaling are the increased angiogenesis and tumor progression [3,20]. PKD-1 is expressed in pNET cells, where it regulates hormone secretion [26] and initiates pancreatic acinar cell reprogramming and progression to intraepithelial neoplasia [27]. PKD-1 signaling is known to promote CSC maintenance in some caners but has not been well explored in pNET CSCs. This pathway promotes stem-like features of cancer cells in estrogen positive breast cancers and regulates endothelial cell (EC) differentiation and arteriolar remodeling [18,21-24,28]. We hypothesized that PKD-1 signaling regulates CSC maintenance in pNETs. Unexpectedly, we identified pNET cells that present stem-like features with cellular plasticity and partial epithelial to mesenchymal transition (EMT). Similar to estrogen-positive breast cancers, pNET CSCs tended to be enriched in the vascular niches. The EMT is known as a cellular transdifferentiation program that enables epithelial cancer cells to become invasive and metastatic [29,30] and stem cell-like [29]. Our study suggests that PKD-1 signaling may endow pNET cells with partial EMT traits and plasticity, enabling them to maintain at flexible CSC states rather than terminal EMT. This may facilitate tumor initiation and completion of the metastatic processes during pNET progression.

## 2. Materials and Methods

### 2.1 Reagents and Antibodies

Oleoyl-L-α-lysophosphatidic acid (LPA, L7260) and a Periodic Acid-Schiff (PAS) Kit (395B) were purchased from Sigma-Aldrich. The PKD inhibitor CRT0066101 (A8679) was purchased from APExBio. The reagents for RT-qPCR include the RNeasy Mini Kit (Qiagen), Power SYBR Green PCR Master Mix and High-Capacity cDNA Reverse Transcription Kits (Applied Biosystems). RT^2^ qPCR Primer assays for GAPDH and primers for target genes were respectively purchased from Qiagen and IDT. Antibodies for immunoblot included PKD/PKCμ (90039S), phospho-PKD/PKCμ (2051T), phospho-Erk (1/2) (4370), Erk (1/2) (137F5), E-cadherin (24E10), vimentin (D21H3) (Cell Signaling Technology) and ALDH1A1 (AF5869) (R&D Systems). Antibodies for immunostaining included rabbit anti-human/mouse PKD-1 (SAB4502371) and mouse anti-human/mouse α-smooth muscle actin (α-SMA, A2547) (Sigma-Aldrich), mouse anti-human/rat CD44 (5640S) and HRP-linked anti-rabbit IgG (7074) (Cell Signaling Technology), CD45 monoclonal Antibody (14-9457-80), AlexaFluor 594 conjugated donkey anti-rabbit IgG (A21207), AlexaFluor 594 conjugated donkey anti-mouse IgG(A21203), AlexaFluor 594 conjugated donkey anti-goat IgG (A11058), AlexaFluor 488 conjugated donkey anti-goat IgG(A11055), AlexaFluor 488 conjugated donkey anti-rabbit IgG (A21206), and AlexaFluor 488 conjugated goat anti-mouse IgG (A11001) (Invitrogen). VECTASHIELD Antifade Mounting Medium with DAPI (H-1200) was purchased from VECTASHIELD. A DAB substrate kit (8059) was purchased from Cell Signaling Technology. Opti-MEM I reduced-serum medium (51985-034) was from Gibco.

### 2.2 Cell Culture

BON and QGP-1 were cultured as described previously [31]. Briefly, cells were grown in glutamine-containing DMEM:F-12 (Gibco) or RPMI1640 (Corning) medium with 5 - 10% fetal bovine serum (FBS) and 1% penicillin/streptomycin under 5% CO_2_ and at 37°C.

### 2.3 Real Time RT-qPCR

mRNA levels were assayed as described previously [32]. Briefly, total RNA was isolated from tumor cells using the RNeasy Mini Kit (Qiagen) and then subjected to cDNA synthesis and RT-qPCR using the CFX Connect Real-Time System (Bio-Rad). The qPCR primers were synthesized by Integrated DNA Technologies IDT, and used in PCR reactions to detect expression of target genes. GAPDH transcripts were amplified in separate wells for normalization, and Qiagen primer assays were also used for GAPDH detection. The relative Ct value was used to compare the fold or quantify changes in mRNA expression.

### 2.4 Immunoblot Assays

Cell lysates were collected by processing with RIPA buffer (Sigma), and protein concentrations were quantified using a BCA kits (Pierce Chemical). Cell lysates were separated on polyacrylamide gel, and proteins transferred to Western blot membranes followed by immunoblotting with appropriate antibodies. NIH Image J (https://imagej.nih.gov/ij/index.html) was used for densitometry to determine relative expression of target proteins.

### 2.5 Human pNET Specimens

Tumor specimens from human patients with pNETs were used to perform immunohistochemistry and immunofluorescence experiments without any link to subject-identifiable information [33]. They included tissue samples from 4 pNET patients, a tissue microarray (TMA) slide generated with pNET tissues samples from 35 patients, and a control TMA slide of human organs from 33 normal individuals (UAB Department of Pathology).

### 2.6 Immunofluorescence and Immunohistochemistry

Tissues were fixed in 10% formalin for paraffin block preparation, sectioned, and processed. Cells were fixed in 4% PFA for immunohistochemical and immunofluorescence staining. Briefly, paraffin tissues were deparaffinized and dehydrated using Histo-Clear and gradient ethanol respectively. Antigen retrieval was performed by immersing slides in antigen unmasking solutions (Vector 21202), placing the slides in a microwave oven for 1 minute and boiling the slides at 95 - 99 °C for 15 min. The slices were cooled down to room temperature and washed with distilled water and PBS. 5% BSA was added to block non-specific reactions. The slices were stained with primary antibodies followed by appropriate secondary antibodies. For immunohistochemistry, the slides were stained with DAB chromogen, hematoxylin, and Periodic Acid-Schiff (PAS) based on the manufacturer’s instructions. The images were captured on an Olympus microscope or an All-in-One Fluorescence Microscope BZ-X810 (Keyence).

Immunofluorescence microscopic analysis for cultured cells was performed as described previously [18,23]. Briefly, cells were fixed with 4% PFA, then permeabilized in 0.1-0.3% Triton-100 in PBS and blocked with 3% BSA. The cells were incubated with appropriate primary antibodies at 4°C overnight, followed by incubation with the appropriate secondary antibodies. The immunofluorescence images were captured on an All-in-One Fluorescence Microscope BZ-X810 (Keyence).

### 2.7 Aldehyde Dehydrogenase (ALDH) Activity Assays

The ALDH1 ELISA kit (ab155894, Abcam) was used to measure ALDH1 activity in BON and QGP-1 cells with different treatments. The reaction mixture of ALDH assay buffer and ALDH substrate was incubated for 20-60 minutes at room temperature. Fluorescence values were recorded at Ex/Em 535/587nm by the Gen5 program (BioTek) and used to calculate ALDH activity based upon the manufacturer’s instruction.

### 2.8 Tumorsphere Formation Assays

2,500 cells were seeded into each well of an ultra-low attached 6-well plate in 2 mL complete MammoCultTM Medium (Stem Cell Technologies) based upon the manufacturer’s instructions. In some experiments, tumorspheres were exposed to vehicle control or reagents every three days. The numbers of tumorspheres were randomly counted with eight repetitions after culture for 7 days. Images of tumorspheres were captured on an OLYMPUS LH50A microscope.

### 2.9 Statistics

Quantitative data are presented as the mean ± SD or SEM as indicated. Data were analyzed by 2-sided unpaired t tests using the GraphPad Prism 9. One-way ANOVA was also used to determine whether there are any statistically significant differences between independent groups.

## 3. Results

### 3.1 Regulation of Stem-like Phenotype in pNETs by PKD-1 Signaling

CSCs are a subpopulation of cells within a tumor that are able to initiate tumors when propagated in animal models, sustain proliferation, and promote metastasis and therapeutic resistance [8,12,34-37]. CSCs and aggressive cancer cells can be found in close proximity to blood vessels as the perivascular niche enables perfusion of oxygen and nutrients [24,38]. To determine the presence and distribution of stem-like cancer cells in pNETs, we stained tissue sections from human pNET patients with the CSC marker CD44, the pericyte/vascular marker smooth muscle-alpha actin (α-SMA), and the lymphatic cell marker CD45. Immunofluorescence microscopy showed that a group of CD44-positive pNET cells were adjacent to vascular networks in the tumor microenvironment. In contrast, the glandular epithelial cells in normal pancreatic tissue expressed minimal levels of CD44 (Figure 1 A & B; Supplementary Figure 1). Similar to our previous study in ER-positive breast cancers [24], a subset of CD44^+^CD45^-^ tumor cells detached from their nests and accumulated near the capillaries or around the α-SMA^+^ arterioles, along with the presence of CD44^+^ and CD45^+^ lymphatic cells (Figure 1C).

**Figure 1.**
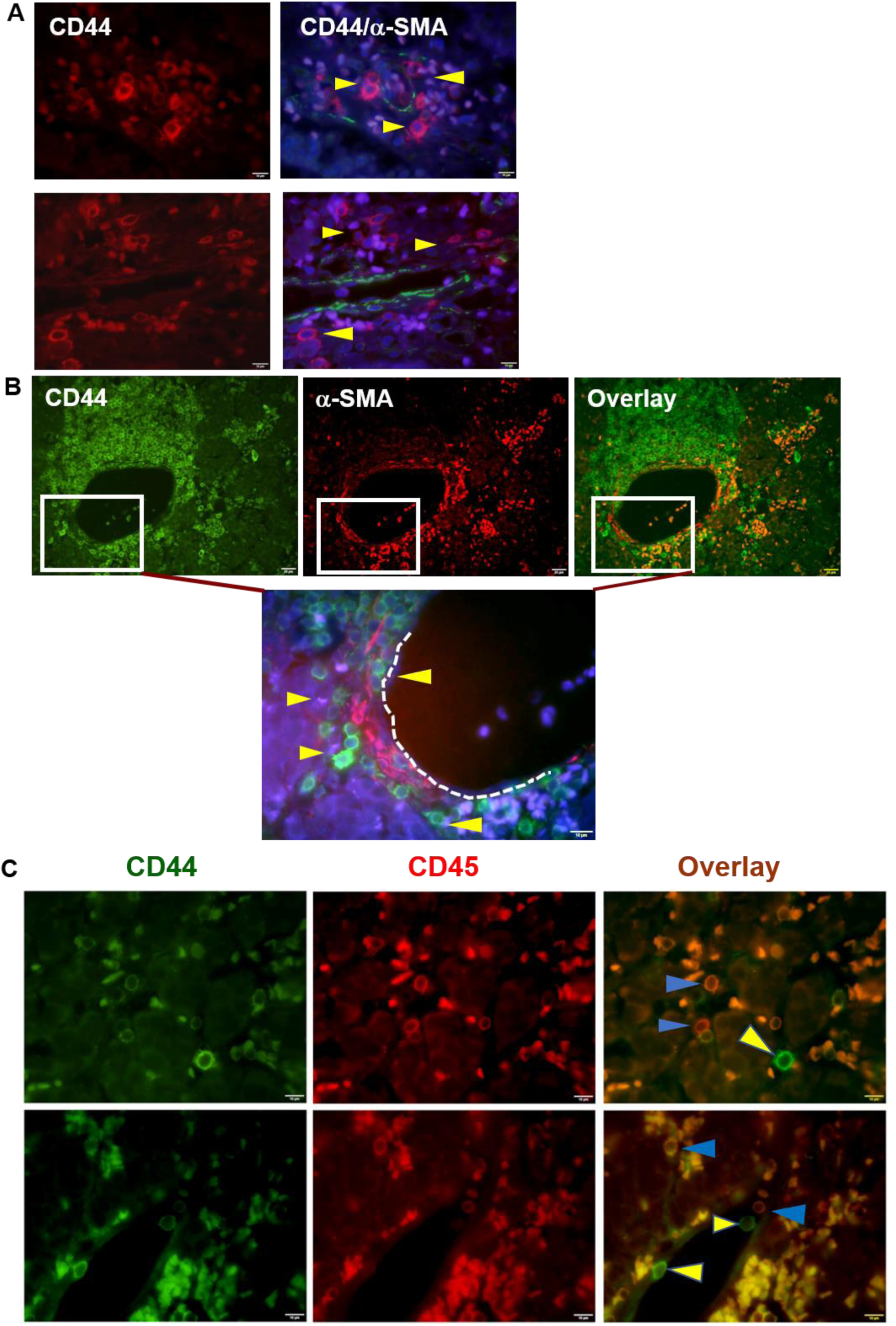
Identification and distribution of CD44-positive cancer stem-like cells in human pNET tissues. **A**. Human pNET specimens in patient 1 were co-stained with CD44 and α-SMA antibodies followed by appropriate secondary antibodies, with DAPI staining the nuclei (blue) by immunofluorescence microscopy. CD44-positive cancer cells (red) marked by yellow arrowheads were close to the α-SMA positive (green) blood vessels. **B**: Human pNET specimens in patient 2 were co-stained as in **A**. CD44-positive cancer cells (green) were indicated by yellow arrowheads, which were close to the α-SMA positive (red) blood vessels. **Note:** A group of CD44-positive cells appear to distribute within the α-SMA-positive vascular niche or nearby blood vessels. Shown are representative images from different individual patients. Bar = 10 or 20 µm. **C**. Human pNET tissues were co-stained with CD44 antibodies and CD45 antibodies followed by appropriate secondary antibodies. Overlay images were collected by immunofluorescence microscopy. CD44-positive (green, yellow arrowheads) and both CD44-positive and CD45-positive cells (orange, blue arrowheads) were observed under a fluorescence microscope. Images were acquired by a fluorescence microscope equipped with a CCD camera. Shown are representative images from two individual patients. Bar = 10 µm. **Note:** Cells positive with both CD44 and CD45 are smaller and regarded as lymphatic cells (orange), a subset of cancer cells are bigger and positive for CD44 but negative for CD45, indicating their stem-like phenotype.

To determine whether PKD-1 pathway is associated with CSCs in pNETs, we examined the expression of PKD-1 and CD44 in tumor tissues from human pNET patients. Interestingly, tumor cells in cancer nests showed low to moderate levels of PKD-1 expression, along with low levels of CD44 expression. In contrast, disseminated tumor cells that were detached from their nests and invaded stroma as well as near blood vessels demonstrated relatively high levels of PKD-1 and CD44. The elevation of PKD-1 and CD44 was particularly high in tumor cells close to the blood vessels (Figure 2A, Supplementary Figure 2). To clarify the role of PKD-1 in the maintenance of a stem cell-like state in pNETs, we genetically targeted PKD-1 in BON cells. As shown in Figure 2B, transfection of siRNA significantly reduced the expression of endogenous PKD-1. Concomitantly, tumorsphere formation capacity was impaired in these tumor cells with PKD-1 knockdown (Figure 2B).

**Figure 2.**
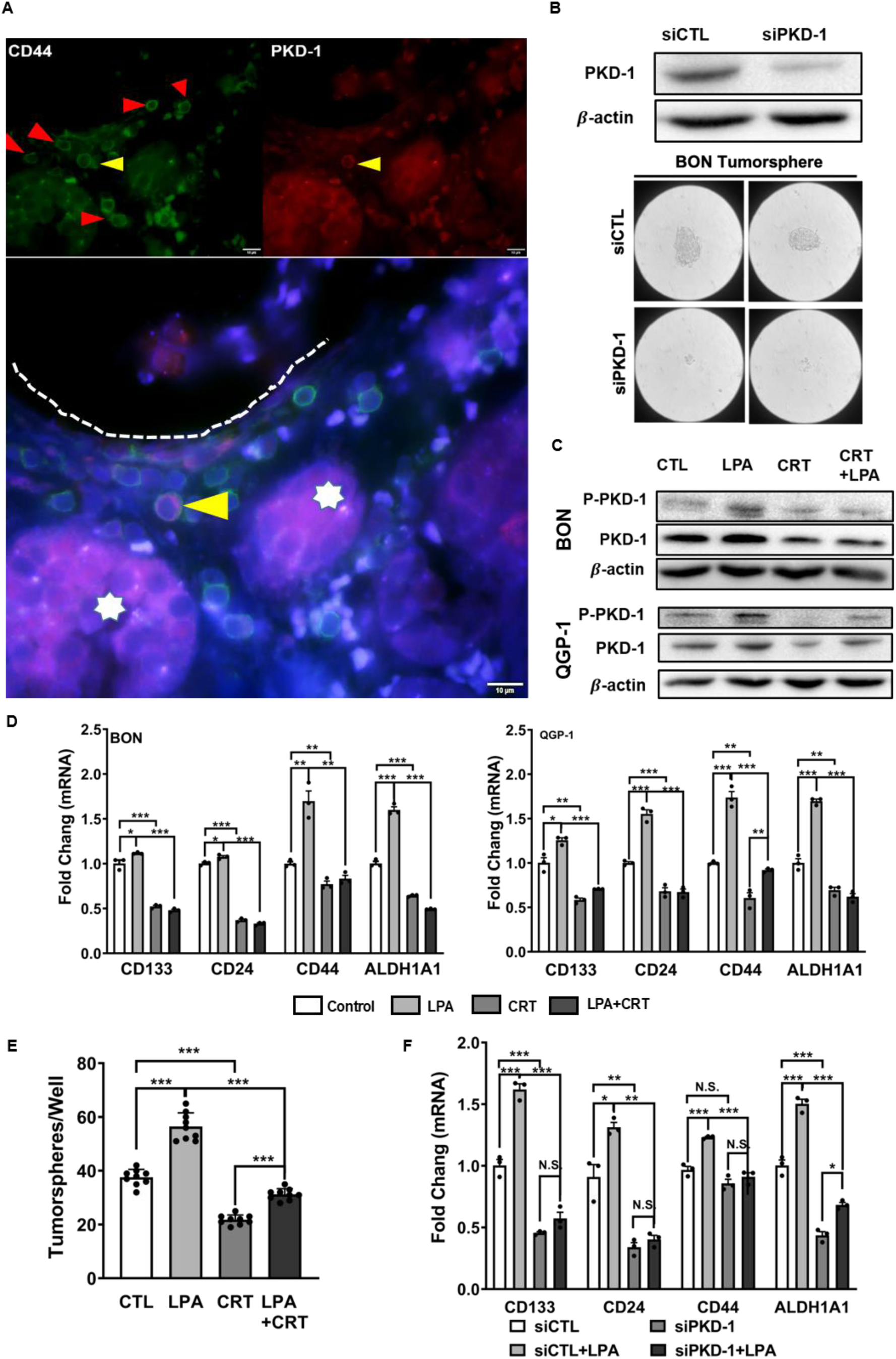
PKD-1 signaling in the maintenance of cancer stem-like features in pNETs. **A**. Distribution of PKD-1^+^ and CD44^+^CSCs within the vascular niche. Human pNET specimens were co-stained with CD44 and PKD-1 antibodies followed by appropriate secondary antibodies, with DAPI staining the nuclei (Blue). Stem-like cells with CD44-positive (green), PKD-1-positive (red) or both positive (pink) were observed under a fluorescence microscope. A few CD44-positive cancer stem-like cells tended to accumulate near the vascular lumen (red arrow heads), and PKD-1-positive CSCs might be leaving tumor nests (stars) for the vascular lumen. The fluorescence images were acquired by an immunofluorescence microscope equipped with a CCD camera. Shown are representative images. Bar = 10 µm. **B**. BON and QGP-1 cells were transfected with siRNA control and siPKD-1 to knock down PKD-1. Knockdown efficiency was confirmed by Western Blots (upper panel). The control and BON cells with knocked-down PKD-1 were subjected to tumorsphere formation assays. Images were acquired by the OLYMPUS CK30 microscope. Representative images are shown for tumorsphere formation (lower panel). **C**. Cell lysates were extracted from BON and QGP-1 cells exposed to the vehicle control, 10 *μ*M LPA, 2 *μ*M CRT0066101, or their combinations after 24 hours. The expression levels of phosphorylated PKD-1 and PKD-1 were detected by Western blots. Shown are representative images. **D**. BON and QGP-1 cells were exposed to 10 *μ*M LPA, 2 *μ*M CRT0066101, or their combination for 24 hours, and total RNA was extracted for the detection of mRNA levels of genes related to stemness properties by RT-qPCR. **E**. Effect of PKD inhibitor in tumorsphere formation. BON cells were cultured in complete MammoCult(tm) medium with the treatment of 10 *μ*M LPA, 2 *μ*M CRT0066101, or their combination for 7 days. The mammary spheres were counted under the OLYMPUS CK30 microscope. **F**. Control and BON cells with PKD-1 knockdown were exposed to 10 *μ*M LPA, 2 *μ*M CRT0066101, or their combination for 24 hours, and total RNA was extracted for the detection of mRNA levels of genes related to stemness properties by RT-qPCR. **Note:** Triplicate experiments were performed, and the results are shown as the mean ± SEM. \**P* < 0.05, \*\**P* < 0.01, \*\*\**P* < 0.001.

Lysophosphatidic acid (LPA), a lipid signaling mediator, activates PKD-1 and promotes tumor initiation, development of cancer stem-like features and metastasis [24,39-43]. To further determine the role of PKD-1 pathway in CSC maintenance, we treated pNET cells with LPA. pNET cells exposed to LPA activated the PKD-1 signaling pathway, and the pathway was effectively targeted with a PKD inhibitor CRT0066101 (Figure 2C, Supplementary Figure 3A & B). Moreover, LPA treatment in the cancer cells stimulated the expression of a CSC-related gene signature via the PKD signaling (Figure 2D), including CD133 and CD44, two common CSC markers whose expression is associated with a poor prognosis in patients with pNETs [44]. Furthermore, pharmacological inhibition of LPA-induced PKD-1 pathway using the PKD inhibitor in pNET cell attenuated tumorsphere formation (Figure 2E), a hallmark of cancer stem-like cells [45]. To further confirm the critical role of PKD-1 signaling in the regulation of stem cell-like features, we knocked down PKD-1 expression in BON cells and examined expression of the stemness-related genes. LPA treatment significantly increased mRNA expression of CD44, CD133, CD24, and ALDH1A1. However, their expression was decreased with PKD-1 knockdown (Figure 2F).

### 3.2 Requirement of PKD-1 Signaling in Partial EMT and CSC Plasticity

EMT programs function as major mechanisms for generating CSCs [29] and tumor invasion and metastasis [30,46-48]. Metastatic tumor cells probably present different epithelial or mesenchymal phenotypes from cells in tumor nests. To determine EMT features in different subsets of tumor cells, we examined the presence of E-cadherin, N-cadherin and vimentin in pNET tissues by immunohistochemistry (IHC), along with Periodic Acid-Schiff (PAS) double-staining for the matrix. We observed that there was expression of the mesenchymal marker vimentin in some tumor cells (Figure 3A & 3B), and these cells tended to be distributed close to the vascular network (Figure 3B) or in a manner similar to PKD-1-positive CSCs (Figure 2A). There was also expression of E-cadherin in most of tumor cells, an indicator of mesenchymal to epithelial reverting transitions during the metastatic seeding of disseminated carcinomas [49], with little expression of the mesenchymal marker, N-cadherin (Figure 3A).

**Figure 3.**
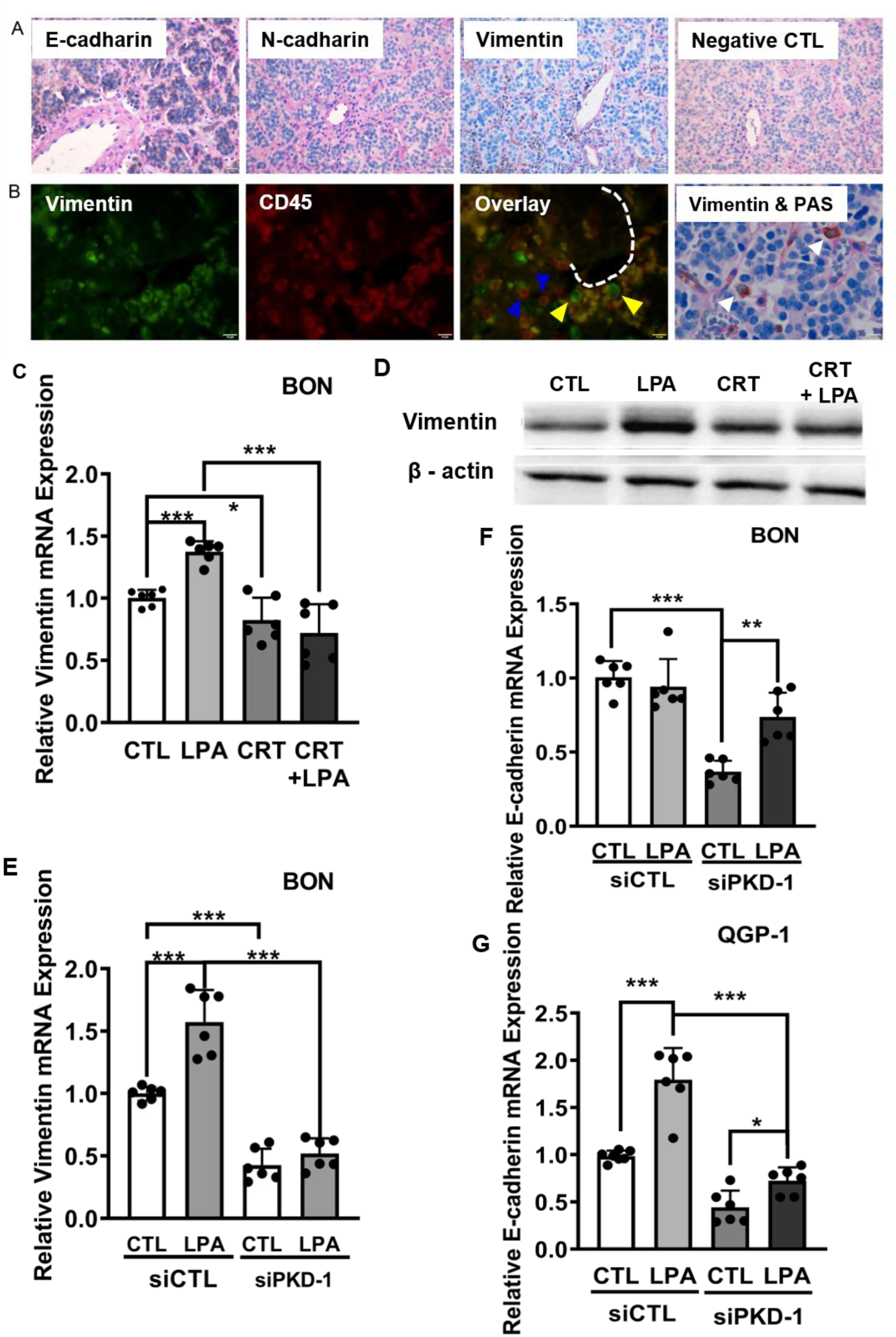
Regulation of EMT by PKD-1 Signaling in pNETs. **A**. pNET tissues from human patients were stained with E-cadherin, N-cadherin and vimentin antibodies by immunohistochemistry (IHC), along with staining by Periodic Acid-Schiff (PAS). Tumor cells expressed E-cadherin but little N-cadherin. A small subset of tumor cells that detached from a cancer nest and distributed in the vascular network expressed mesenchymal marker vimentin. Bar = 20 µm. **B**. pNET tissues from human patients were co-stained by vimentin and CD45 antibodies followed by appropriate secondary antibodies. Vimentin-positive (green) and/or CD45-positive (red) cells were observed under an immunofluorescence microscope. The fluorescence images were acquired by an immunofluorescence microscope equipped with a CCD camera. Shown are representative images. Bar = 10 µm. **Note:** Vimentin-positive CD45-negative tumor cells (yellow arrowhead), that mainly located within the vascular network, from vimentin-positive CD45-positive lymphatic cells (blue arrowhead). Double-staining with IHC and PAS showed that these vimentin-positive mesenchymal tumor cells (white arrowhead) are located in the vascular network or detached from their nests, and exhibited larger nuclei compared to lymphatic cells (lower panel, Vimentin & PAS). Antigens are shown as brown color by IHC (HRP-DAB), and vascular basement membrane was shown as pink by PAS-staining. Bar = 10 µm. **C**. BON cells were treated with 10 *μ*M LPA, 2 *μ*M CRT0066101, or their combination in serum-free medium for 24 hours. Total RNA was extracted from each group to assay mRNA levels of vimentin by RT-qPCR. **D**. BON cells were cultured in DMEM/F12 medium with 5% FBS. After starvation in serum-free DMEM/F12 medium for 6 hours, the cells were treated with 10 µM of LPA, and/or 5 µM of CRT in serum-free DMEM/F12 medium for 24 hours under 5% CO2 at 37°C. Cell lysates were collected and subjected to Western blots for vimentin expression. Shown is a representative image. **E**. BON cells were transfected with scramble control or PKD-1 siRNA for 24 hours, followed by treatment with 10 *μ*M LPA for 24 hours. Total RNA was isolated for vimentin gene expression by RT-qPCR. **F**. BON cells were transfected and treated as **E** for E-cadherin gene expression by RT-qPCR. **G**. QGP-1 cells were transfected and treated as **E** for E-cadherin gene expression by RT-qPCR. **Note:** Triplicate experiments were performed, and the results are shown as the mean value ± SEM. \**P*<0.05, \*\**P*<0.01, \*\*\**P*<0.001.

LPA is known to regulate an EMT program [50]. To determine whether PKD-1 signaling regulates vimentin expression, we exposed pNET cells to LPA, the PKD inhibitor or their combination and examined vimentin mRNA levels in response to LPA/PKD-1 signaling. Intriguingly, exposure to LPA increased expression of vimentin mRNA in BON (Figure 3C) and QGP-1 cells (Supplementary Figure 4A). Treatment with a PKD inhibitor prevented LPA-mediated induction of vimentin at mRNA levels (Figure 3C & Supplementary Figure 4A). Immunoblotting confirmed that the PKD inhibitor repressed LPA-induced vimentin protein expression (Figure 3D). Furthermore, PKD-1 knockdown reduced endogenous vimentin expression and abolished LPA-mediated upregulation of vimentin mRNA in BON (Figure 3E) and in QGP-1 cells (Supplementary Figure 4B). LPA treatment in BON cells did not increase E-cadherin mRNA but PKD-1 knockdown downregulated endogenous expression, which was rescued by treatment with LPA (Figure 3F). Similar to BON cells, genetic targeting of PKD-1 in QGP-1 cells decreased the expression of endogenous E-cadherin. However, LPA exposure was able to increase the expression of E-cadherin in QGP-1 cells, which was attenuated by knocking down PKD-1 (Figure 3G).

ALDH1 activity is essential for CSC plasticity and metastatic potential [51,52]. Given that PKD-1 signaling plays a pivotal role in the regulation of ALDH1A1 in pNET CSCs (Figure 2C & D), we treated BON cells with LPA, a PKD inhibitor or their combination, and examined ALDH1A1 expression. We did not observe significant changes in expression of ALDH1A1 protein by immunoblotting in BON cells exposed to LPA and/or a PKD inhibitor (data not shown). However, immunofluorescence demonstrated that the percentage of cells with enhanced ALDH1A1 expression (ALDH1A1^+^) was increased with LPA treatment (P<0.01), and this effect was prevented by co-treatment with a PKD inhibitor (P<0.05) (Figure 4A). To validate the essential role of PKD-1 signaling in ALDH1A1 levels, we knocked down endogenous PKD-1 expression in pNET cells. Knocking down PKD-1 led to a decrease in the number of ALDH1A1^+^ BON cells (Figure 4B). The number of cells with enhanced ALDH1A1 expression was also reduced in those QGP-1 cells with knocking down PKD-1 when compared to the control (Supplementary Figure 5A). Furthermore, LPA treatment moderately increased ALDH1 activity, and this increase was attenuated by co-treatment with a PKD inhibitor in both BON cells (Figure 4C) and QGP-1 cells (Supplementary Figure 5B) by co-treatment with a PKD inhibitor. Finally, PKD-1 knockdown decreased ALDH1 activity in BON cells when compared with the scrambled control (Figure 4D). Together, the results indicate that PKD-1 signaling may promote CSCs characterized by plasticity and partial EMT, likely enhancing metastatic potential in pNET cells.

**Figure 4.**
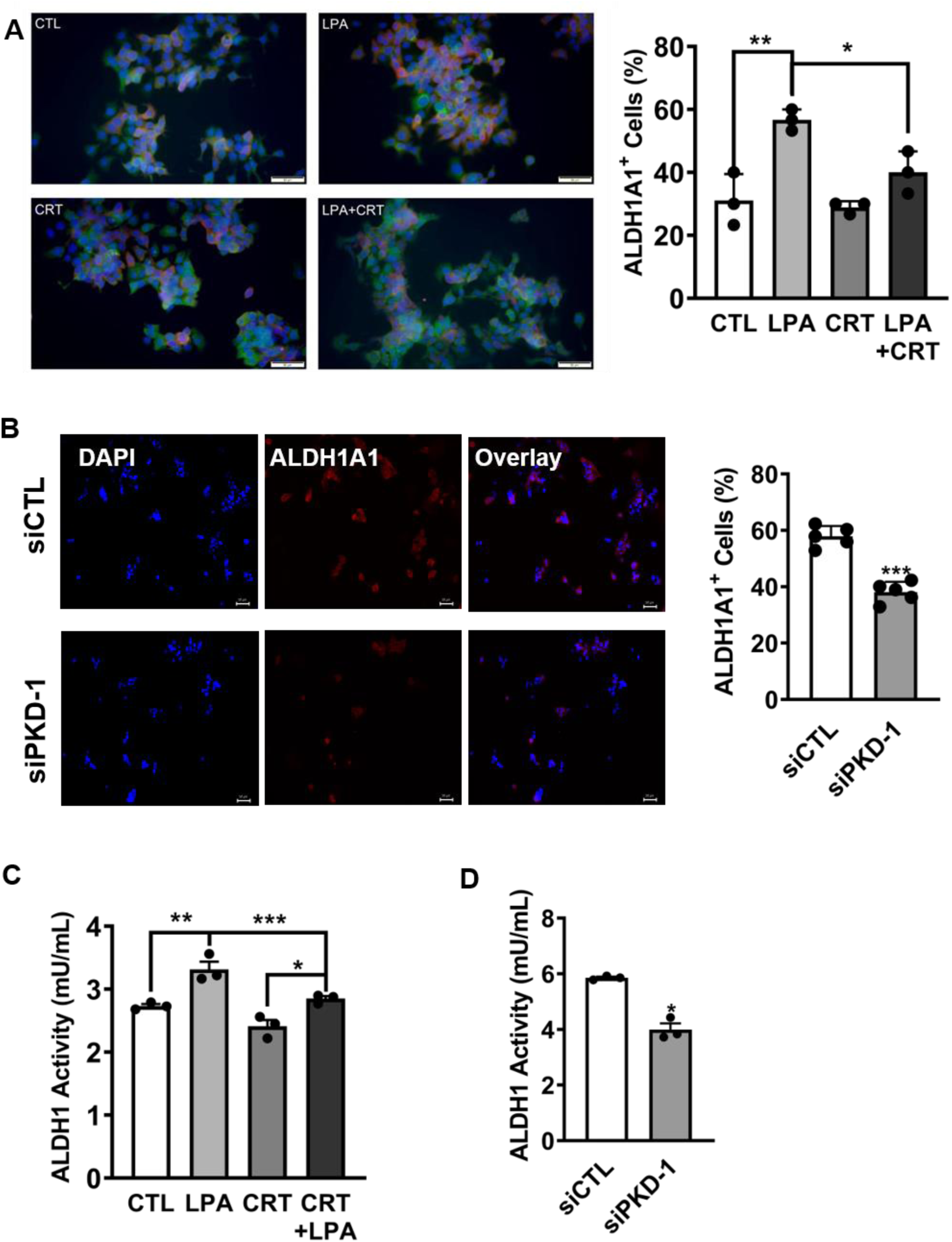
Association of PKD-1 signaling with ALDH1. **A**. BON cells were cultured in DMEM/F12 medium with 5% FBS. After starvation in serum-free DMEM/F12 medium for 6 hours, the cells were treated with 10 µM of LPA, and/or 5 µM of CRT in serum free DMEM/F12 medium for 24 hours under 5% CO2 at 37°C. The cells were incubated with ALDH1A1 and PKD-1 antibodies followed by appropriate secondary antibodies. The percentages of cells with high levels of ALDH1A1 expression (red) were counted randomly up to 30 individual cells and triple counting was performed. Graphpad Prism 9 was used for statistical analysis. **B**. BON cells were transfected with scramble control or PKD-1 siRNA for 24 hours, and the cells were processed for staining with an ALDH1A1 antibody followed by an appropriate secondary antibody. The cells with high levels of ALDH1A1 expression were observed and counted under a fluorescence microscope by counting up to 100 cells randomly in each field. Five repetitions were performed, and the statistic difference was calculated by GraphPad Prism 9. **C**. BON cells were treated with 10 *μ*M LPA, 2 *μ*M CRT0066101, or their combination for 24 hours. ALDH1 activity was measured by ELISA in a plate reader. **D**. BON cells were transfected with scramble control or PKD-1 siRNA to knock down endogenous expression. ALDH1 activities were measured by ELISA in a plate reader. Triplicate experiments were performed. The results were shown as the mean ± SEM. \**P* < 0.05, \*\**P* < 0.01, \*\*\**P* < 0.001.

### 3.3. Critical Role of PKD-1 Signaling in CD36 Expression in pNET Cells

CD36 is known as a scavenge receptor, fatty acid receptor and angiogenesis regulator [18,21,23,24,53,54]. The association of CD36 with tumorigenesis is controversial [23,24,54-57]. However, recent studies demonstrated that CD36 drives the CSC phenotype, and increases drug resistance capacity and metastatic potential of CSCs [24,54,57,58]. In pancreatic cancers, the expression of CD36 may be negatively associated with the progression of pancreatic adenocarcinoma [59]. Since PKD-1 signaling downregulates CD36 expression in vascular ECs [18,21,24], we intended to determine whether PKD-1 signaling regulates CD36 expression in pNETs. Toward this end, BON and QGP-1 cells were exposed to LPA and/or a PKD inhibitor. Unexpectedly, LPA treatment increased the expression of CD36 at the levels of transcription and translation in both BON (Figure 5A) and QGP-1 cells (Figure 5B). Addition of a pharmacological PKD inhibitor prevented LPA-induced CD36 expression (Figure 5A & B). To validate the essential role of PKD-1 signaling in CD36 expression, we genetically knocked down endogenous expression of PKD-1. Compared with the control group, LPA stimulated mRNA expression of CD36 in BON cells whereas PKD-1 knockdown prevented the LPA-induced expression (Figure 5C). Genetic targeting of PKD-1 also decreased endogenous expression of CD36 at the transcriptional (Figure 5D) and translational levels (Figure 5E) in both BON and QGP-1 cells. The results cherish an important role of PKD-1 signaling in driving CD36 expression and suggest a role of PKD-1 in enhancing metastatic potential and drug resistance in pNETs via CD36-mediated fatty acid metabolism [57,58].

**Figure 5.**
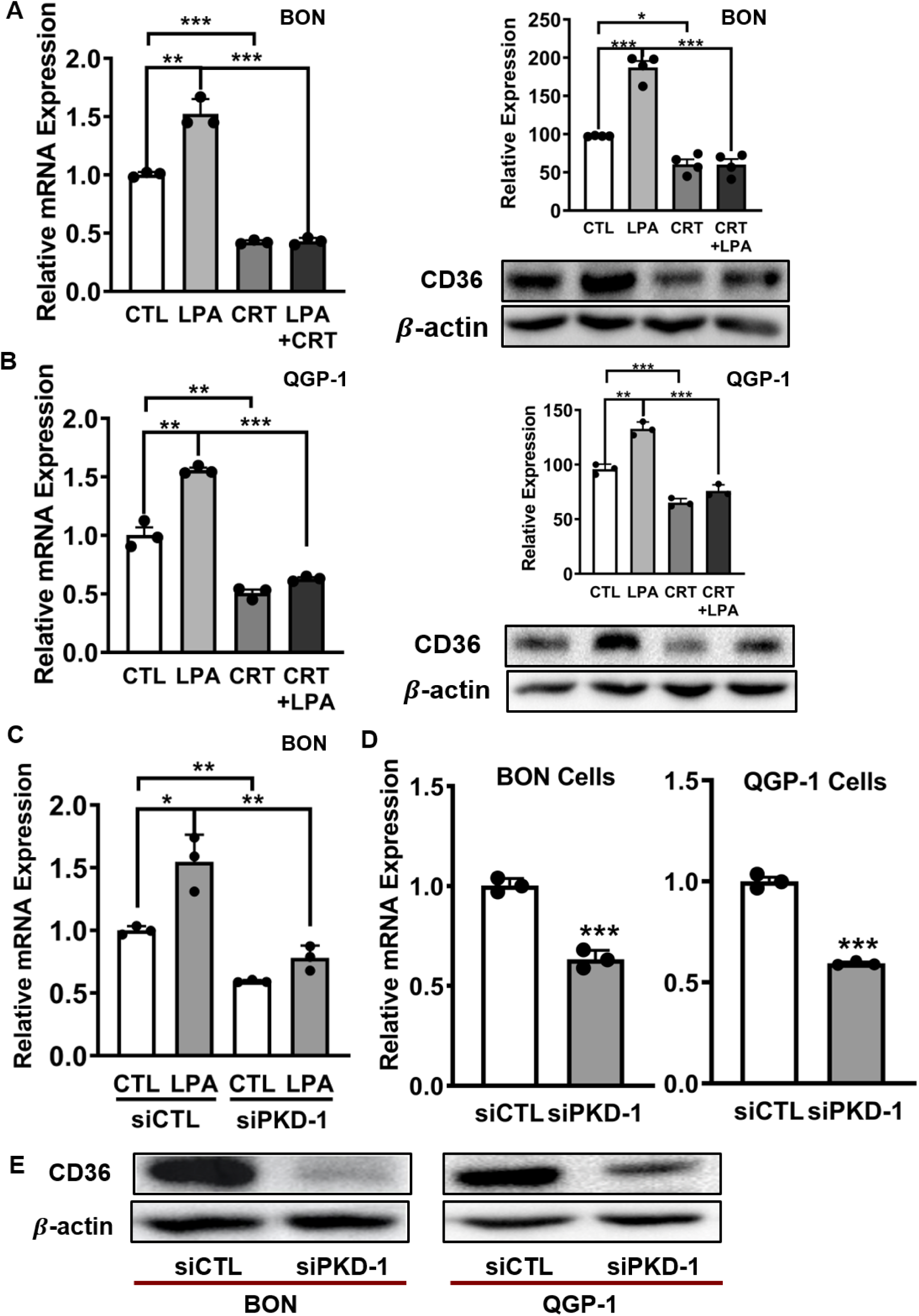
Regulation of CD36 expression by PKD-1 signaling in pNET cells. **A**. BON cells were exposed to 10 *μ*M LPA, 2 *μ*M CRT0066101, or their combination for 24 hours. Total RNA was isolated for CD36 gene expression by RT-qPCR (left panel), and cell lysates were collected and subjected to Western blot for CD36 protein expression (right panel). **B**. QGP-1 cells were treated and assayed for CD36 gene and protein expression as **A. C**. BON cells were transfected with scramble control or PKD-1 siRNA for 24 hours, followed by treatment with 10 *μ*M LPA for 24 hours. Total RNA was isolated for CD36 gene expression by RT-qPCR. **D**. pNET cells were transfected with scramble control or PKD-1 siRNA to knock down endogenous PKD-1 gene expression, and total RNA was isolated for CD36 gene expression by RT-qPCR. **E**. pNET cells were transfected with scramble control or PKD-1 siRNA to knock down endogenous PKD-1 gene expression, and cell lysates were collected and subjected to Western blots for CD36 protein levels. Shown are representative images. **Note:** CD36 protein levels were assessed by densitometry with NIH Image J. Triplicate experiments were performed, and the results are shown as the mean ± SEM. \**P*<0.05, \*\**P*<0.01, \*\*\**P*<0.001.

## 4. Discussion

CSCs that diverge in gene expression are responsible for tumor heterogeneity drive metastasis and therapeutic resistance in a variety of cancers [8,11,12,24,54,57]. These cancer stem-like cells also present in heterogeneous pNETs [6,13,14,37,60]. Our findings provide new insight into the mechanisms and potential roles of PKD-1 signaling in the maintenance of CSCs in pNETs. First, we demonstrate that previously uncharacterized CSCs that are positive for PKD-1 and CD44 are present in human pNETs. They appear to detach from their nests and accumulate within the vascular niches, suggesting the potential of invasion and metastasis due to their stem-like features. Furthermore, PKD-1 signaling is required for the maintenance of unique CSCs with plasticity and partial EMT by regulation of a potential CSC gene signature characterized by stem cell-like plasticity [14,24,54,57,61,62] and expression of both epithelial and mesenchymal markers in pNET cells.

Many signaling pathways associated with oncogenesis including the Notch, Sonic hedgehog and Wnt pathways regulate self-renewal of CSCs [8,29,63,64], and distinct pathways control CSC self-renewal in different tissues. This study highlights a new role of PKD-1 signaling in the regulation of cancer stemness-like features with partial EMT and epithelial/mesenchymal plasticity in pNETs. This is consistent with studies intimating that EMT confers tumor-initiating and metastatic potential to cancer cells, thereby generating high-grade invasive cells with stem cell–like features [29,65,66]. EMT may be a key step in tumorigenesis in pNETs [67]. However, our study demonstrates that in human pNET tissues E-cadherin is constitutively expressed, along with vimentin expression in some cells. In pNET CSCs PKD-1 signaling likely contributes to the establishment of a partial EMT program by induction of vimentin and E-cadherin expression. This phenotype is supported by the fact that MAPK/ERK and PI3K/Akt pathways, an upstream of PKD-1 signaling, interact with a series of intracellular signaling networks to determine the actual implementation of the EMT program at cellular levels [30,68].

E-cadherin is known to be generated in most differentiated tumors [68]. Loss of E-cadherin expression appears to be significantly involved in EMT and suppression of tumor invasion [68,69]. However, it is reasonable to speculate that these pNET cells can maintain an invasive phenotype as high levels of vimentin, ALDH1A1 and CD36 may counteract invasion-suppressor role of E-cadherin. Meanwhile the hybrid states of CSCs facilitate collective cell migration by providing a “stemness window” rather than completely committed toward the mesenchymal phenotype [70]. We hypothesize that by maintaining a partial EMT phenotype that lies between the epithelial and the mesenchymal state, these cells can thus show better CSC plasticity, and may increase metastatic potential due to CD36-mediated fatty acid metabolism [24,52,57,71,72]. The hybrid cell state with mixed epithelial-mesenchymal phenotypes with retention of certain epithelial traits may be central to functionality of CSCs in pNET progression and acquirement of more invasive capacities. This PKD-1 signaling-mediated phenotype in pNETs is different from other types of tumors, where PKD-1 signaling was considered to maintain the epithelial phenotype [73], and warrants further investigation.

Furthermore, this study suggests that in the initial stage PKD-1 signaling might promote the clustering of E-cadherin in pNET cells. This is also different from other cancer types where E-cadherin is typically repressed during EMT, and those cancer cells cannot undergo collective movement due to E-cadherin deficiency [29]. Conversely, the E-cadherin-mediated clusters of cells in pNETs may undergo malignant progression and collective dissemination [74-76] due to the concurrent expression of both E-cadherin and vimentin. This behavior can be further strengthened following concomitantly induced expression of ALDH1A1 and CD36 in response to such environmental factors as LPA that exists in the tumor microenvironment [21]. Furthermore, these factors may regulate tumor cell-microenvironment interaction, thereby promoting partial EMT and CSC maintenance [24,40,42,50]. The E-cadherin-positive cells could easily revert to the epithelial state during metastatic dissemination by undergoing mesenchymal-epithelial reverting transitions (MErT) likely due to their cellular plasticity [49], likely enabling pNTE cells to establish secondary colonies in the liver, the most frequent organ for pNET spread. By further stimulating stemness-like features, the PKD-1 signaling pathway might contribute significantly to aggressive and metastatic behavior in pNETs. Finally, similar to our recent study [24], pNET CSCs could move toward the vascular niches, in which vascular ECs are a key player. Given the critical role of ECs in arteriolar differentiation and tumor progression [8,10,24,77-79], ECs in highly vascularized pNETs may nurture CSCs by direct interactions and indirect generation of such vascular niche factors as LPA, likely via communication with the PKD-1 [6,8,18,24], thereby leading to progression toward malignancy, drug resistance and metastasis. (Figure 6).

**Figure 6.**
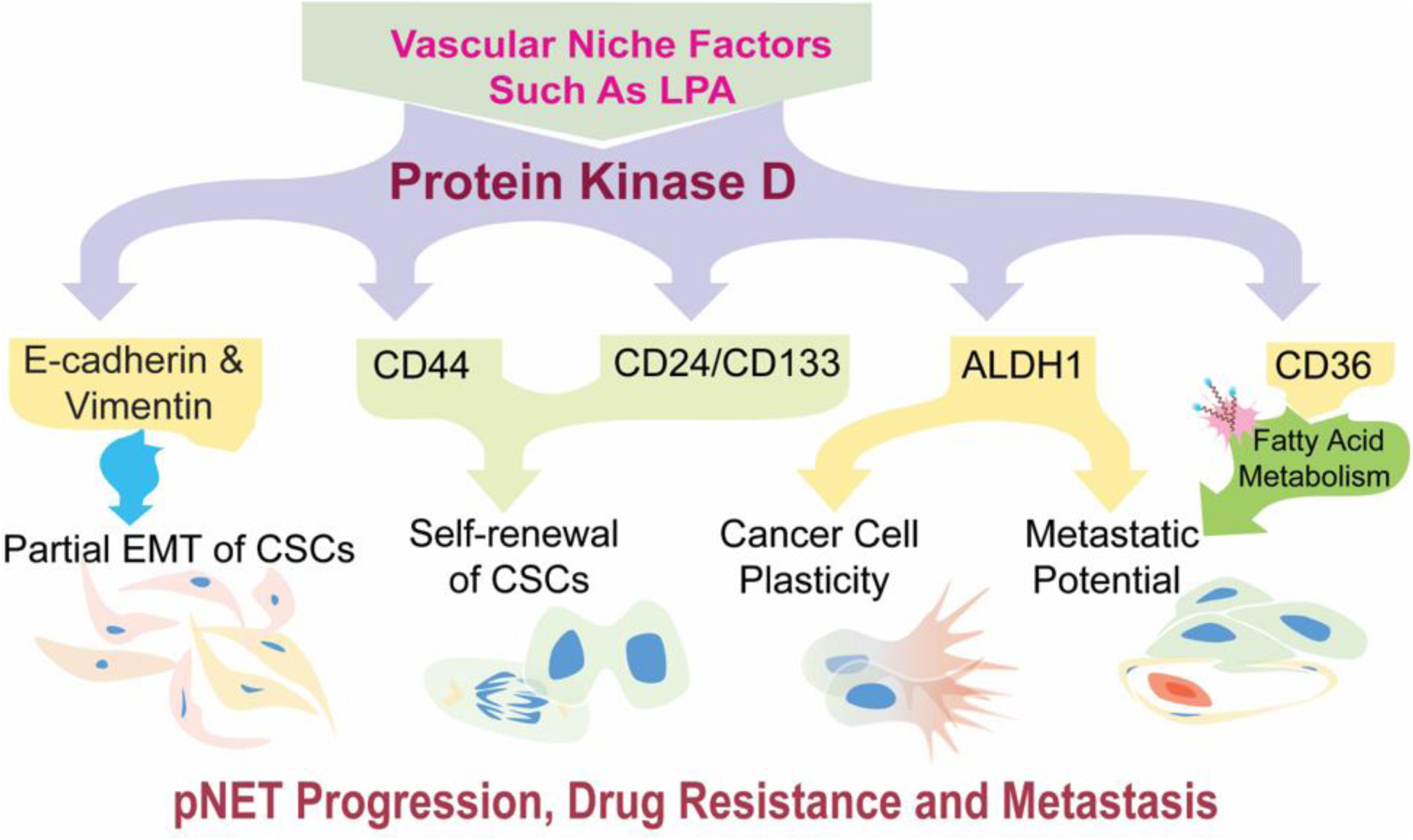
Working model: Regulation of CSCs with plastic and partial EMT phenotype by PKD-1 signaling in pNETs. PKD-1 signaling may induce intermediate or partial EMT with a plastic phenotype in cancer stem-like cells, thereby endowing pNETs with robust invasive and pro-metastatic traits and drug resistance. Vascular niche factors may contribute to this phenotype.

In summary, PKD-1 signaling is required for the maintenance of CSCs with EMT plasticity in pNETs by activation of a partial EMT program and induction of ALDH1 and CD36 expression. This PKD-1 signaling-mediated CSC phenotype might uniquely contribute to the secondary colonization of metastatic cancer cells in the liver. As CD36 expression in cancer cells drives stem cell-like traits and promotes drug resistance and metastatic potential of CSCs [24,54,57,58], PKD-1-mediated CD36 expression may play an important role in pNETs, very likely via CD36-mediated fatty acid metabolism. This concept deserves further investigation and characterization. Although PKD-1 is considered to maintain an epithelial phenotype by negatively regulating key molecules that control EMTs in some cancer cells [73], this study demonstrates that PKD-1 signaling in pNETs is required for concomitant expression of vimentin and E-cadherin in CSCs. This may override the role of PKD-1 in promoting the epithelial phenotype [80], likely leading to malignant progression and metastasis by activation of a partial EMT program and maintenance of CSC traits. This unique phenotype may be explained by the fact that EMT is considered a reversible process that transiently places epithelial cells into mesenchymal cell states [7, 8, 9]. The observed partial EMT status and increased expression of ALDH1A1 may render pNET cells more plastic [24,52,71], thereby conferring high metastatic potential to the CSCs by acquiring migratory and invasive properties.

In addition, not only does PKD-1 signaling promote metastatic potential by upregulating expression of vimentin and increasing the number of ALDH1^+^CSCs [81-83], this signaling also increases the expression of CD36 to activate fatty acid metabolism and further enhances metastatic potential [57]. Therefore, identification of the PKD-1 pathway in CSC plasticity with a partial EMT phenotype may provide new insight into CSC biology in a variety of cancer types since EMT is important during the metastatic stage while E-cadherin-induced MET will aid in the subsequent colonization [29,30]. The hybrid cell states with mixed epithelial-mesenchymal phenotypes may benefit the maintenance of stemness traits in pNETs [70]. It would be interesting and of significance to further investigate precise mechanisms in animal models and the clinic settings. Finally, CSC-EC interactions within the vascular niches in pNET progression merits further characterization since CSCs tend to be enriched within the vascular niches. These additional studies will facilitate the discovery of potential therapeutic interventions and determine possible biomarkers in the prediction of an unfavorable prognosis and relapse in patients with pNETs [12,44,67,84].

## 5. Conclusions

PKD-1 signaling is important in the maintenance of a subpopulation of CSCs characterized by cellular plasticity and partial EMT in pNETs. By regulation of the expression of a cancer stemness-related gene signature, which includes CD36, ALDH1, and vimentin and E-cadherin during EMT/MET processes, this pathway may promote malignant progression, metabolic reprogramming, drug-resistance, and metastasis in pNETs. The study provides insight into the understanding of CSC-mediated tumor progression and offers means of development of potential new therapeutic strategies in a variety of highly angiogenic cancers.

## Supplementary Materials

The following are available online at …… **Figure S1-5**. ……

## Author Contributions

YG and YJ performed experiments and analyzed the data, provided initial figures and figure legends, and contributed substantially to the writing of the manuscript. JBR, RJ, ABH, and HC edited and reviewed the manuscript. BR conceptualized the studies, interpreted the data and wrote the manuscript.

## Acknowledgement

We appreciate the help from Dr. Sameer AL Diffalha in the Department of Pathology, UAB Heersink School of Medicine.

## Supplementary Figures

**Supplementary Figure 1.**
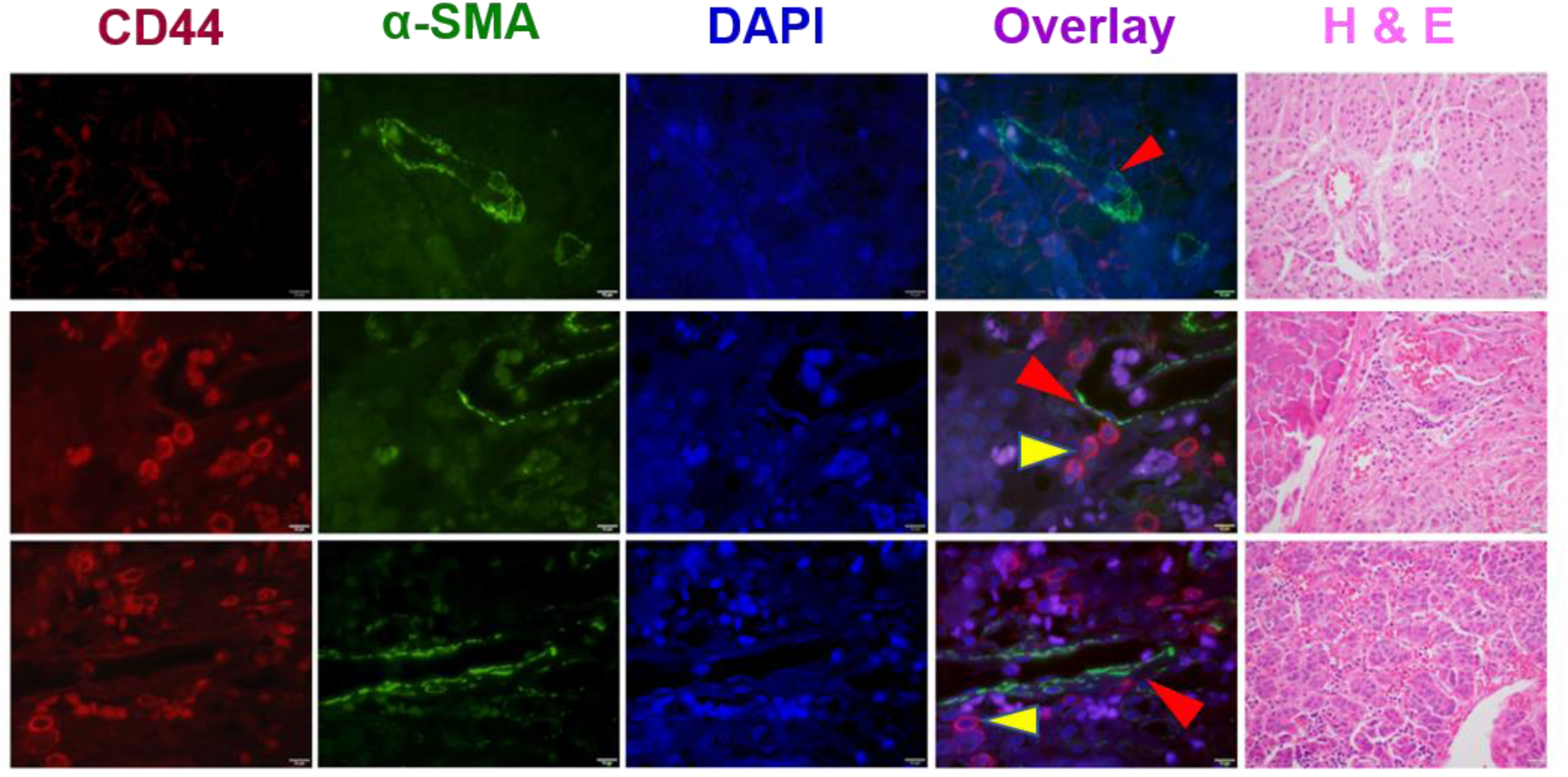
Stem-like phenotype of cancer cells in pNETs. **A**. A human pancreatic tissue control (top panel) and human pNET specimens from two individual patients (middle and lower panels) were co-stained with CD44 and α-SMA antibodies followed by appropriate secondary antibodies, with DAPI staining the nuclei (blue) by immunofluorescence microscopy. CD44-positive cancer stem-like cells were marked by yellow arrowheads, which were close to the α-SMA positive (green, red arrowheads) blood vessels. The fluorescence images were acquired by an immunofluorescence microscope equipped with a CCD camera, and representative images are shown, along with H & E staining for tumor tissue structures. Bar = 10 µm.

**Supplementary Figure 2.**
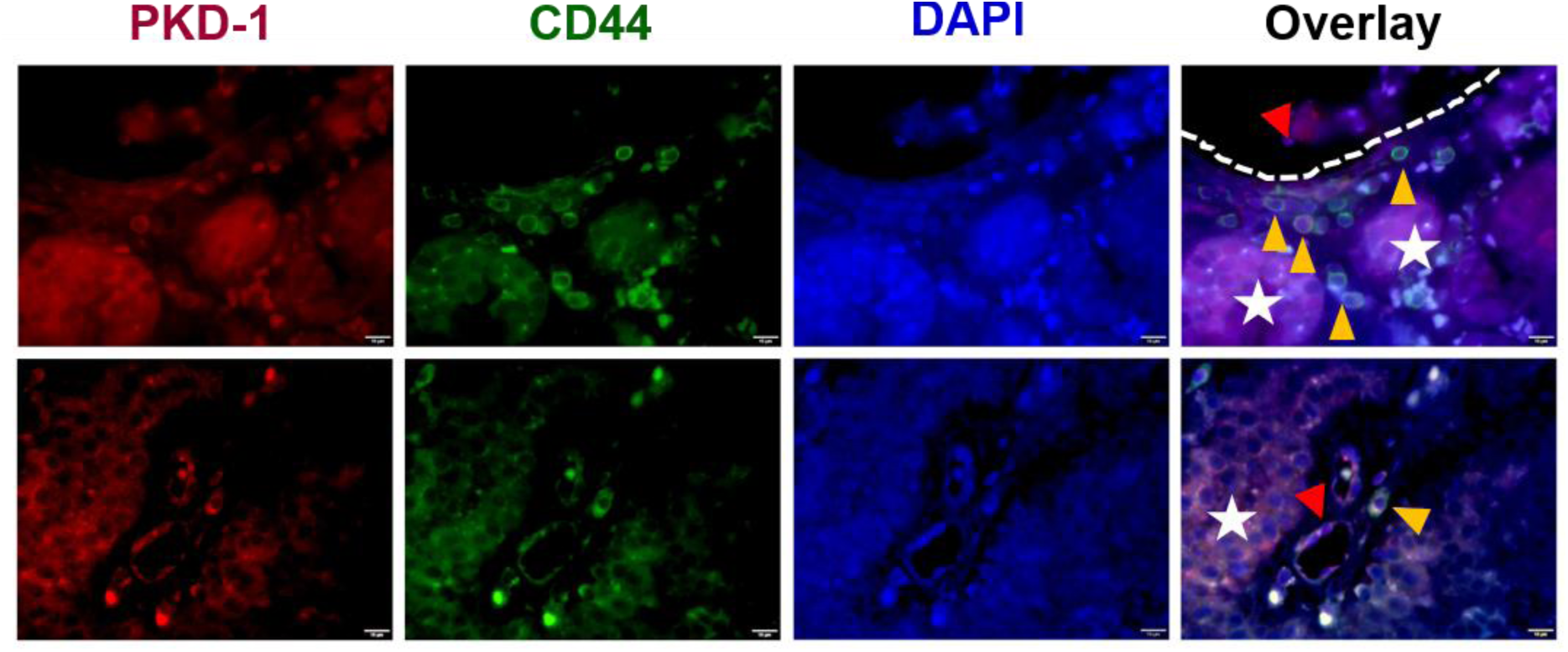
Distribution of CD44-and/or PKD-1-positive cancer stem-like cells in pNET tissues. Human pNET specimens were co-stained with CD44 antibodies and PKD-1 antibodies followed by appropriate secondary antibodies, with DAPI staining the nuclei (blue). CD44-positive (green), PKD-1-positive (red) or both positive were observed under a fluorescence microscope. Cancer cells with high levels of both CD44 and PKD-1 (yellow arrowheads) likely left the tumor nests (white stars) and accumulated in the nearby vascular lumen (red arrowheads). Fluorescence images were acquired by an immunofluorescence microscope equipped with a CCD camera. Shown are representative images from two individual patients. Bar = 10 µm. **Note:** These are additional pictures for Figure 2A.

**Supplementary Figure 3.**
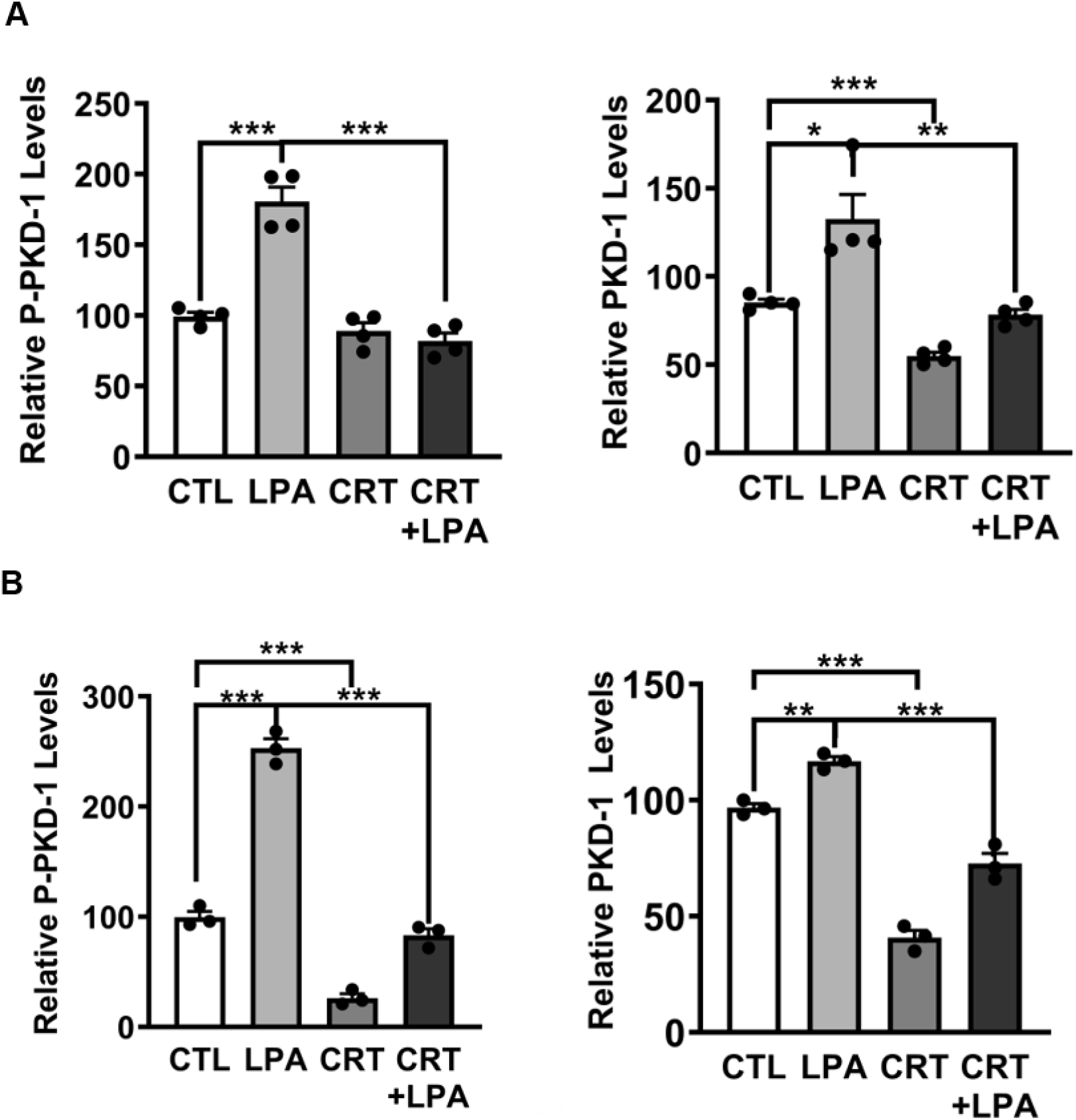
LPA activated PKD-1 signaling pathway in pNET cells. BON cells (**A)** and QGP-1 cells (**B**) were exposed to the vehicle control, 10 *μ*M LPA, 2 *μ*M CRT0066101, or their combinations after 24 hours. Cell lysates were extracted and the expressions of phosphorylated and total PKD-1 were detected by Western blots. Shown are the relative expression levels assessed by densitometry using an NIH Image J. Triplicate experiments were performed, and the results are shown as the mean ± SEM. \**P* < 0.05, \*\**P* < 0.01, \*\*\**P* < 0.001.

**Supplementary Figure 4.**
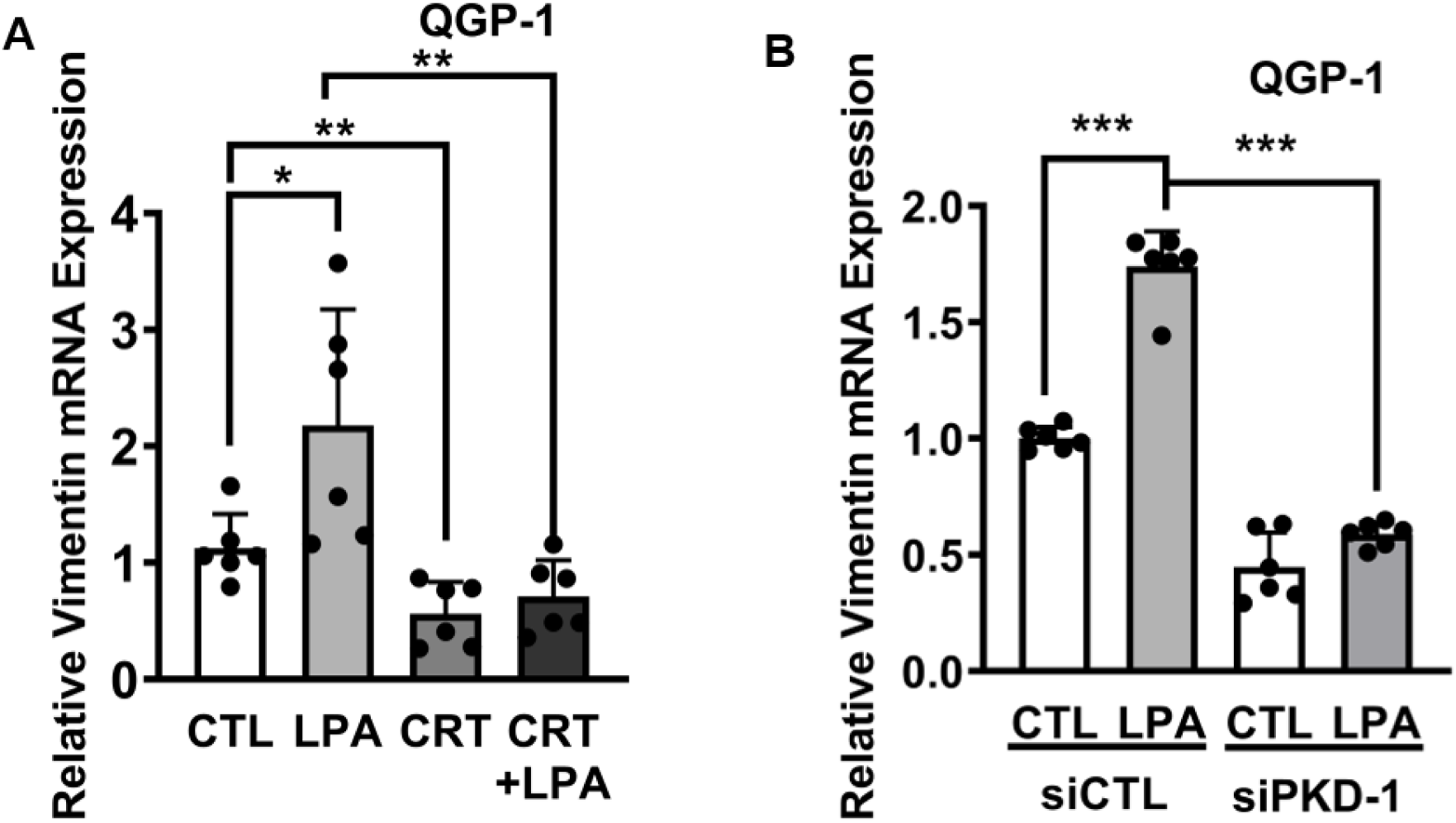
Critical role of PKD-1 signaling in the expression of vimentin in pNET cells. **A**. QGP-1 cells were treated with 10 *μ*M LPA, 2 *μ*M CRT0066101, or their combination in serum-free medium for 24 hours. Total RNA were isolated from each group to detect mRNA levels of vimentin by RT-qPCR. **B**. QGP-1 cells were transfected with scramble control or PKD-1 siRNA for 24 hours, followed by treatment with 10 *μ*M LPA for 24 hours. Total RNA was isolated for vimentin gene expression by RT-qPCR. Triplicate experiments were performed, and the results are shown as the mean ± SEM. \**P* < 0.05, \*\**P* < 0.01, \*\*\**P* < 0.001.

**Supplementary Figure 5.**
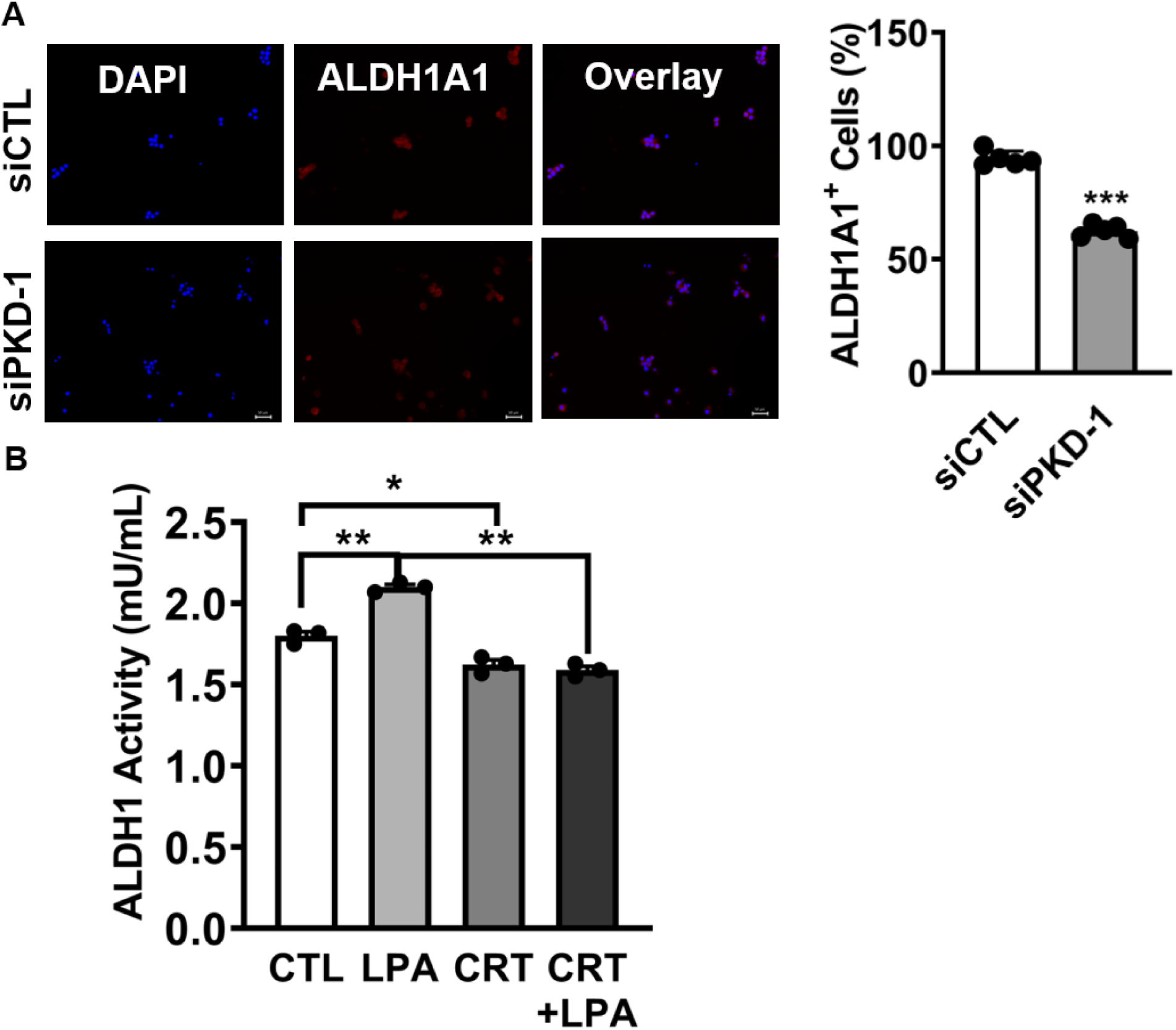
Critical role of PKD-1 signaling in the regulation of ALDH1 expression and activity in pNET cells. **A**. QGP-1 cells were transfected with scramble control or PKD-1 siRNA for 24 hours. Cells were stained with ALDH1A1 antibodies followed by appropriate secondary antibodies. The cells with high levels of ALDH1A1 expression were observed and counted under a fluorescence microscope by counting up to 100 cells randomly in each field. Fluorescence images were acquired by an immunofluorescence microscope equipped with a CCD camera, and representative images are shown. Five repetitions were performed and the statistic difference was calculated using GraphPad Prism 9. \*\*\**P* < 0.001. **B**. QGP-1cells were treated with 10 *μ*M LPA, 2 *μ*M CRT0066101, or their combination for 24 hours. ALDH1 activity was measured by ELISA in a plate reader. \**P* < 0.05, \*\**P* < 0.01.

